# Local Field Potentials, Spiking Activity, and Receptive Fields in Human Visual Cortex

**DOI:** 10.1101/2022.08.28.505627

**Authors:** Lu Luo, Xiongfei Wang, Junshi Lu, Guanpeng Chen, Guoming Luan, Wu Li, Qian Wang, Fang Fang

**Author notes:** **Corresponding Authors:** Fang Fang, Qian Wang.

## Abstract

The concept of receptive field (RF) is central to sensory neuroscience. Neuronal RF properties have been substantially studied in animals, while those in humans remain nearly unexplored. Here, we measured neuronal RFs with intracranial local field potentials (LFPs) and spiking activity in human visual cortex (V1/V2/V3). We recorded LFPs via macro-contacts and discovered that RF sizes estimated from low-frequency activity (LFA, 0.5 – 30 Hz) were larger than those estimated from low-gamma activity (LGA, 30 – 60 Hz) and high-gamma activity (HGA, 60 – 150 Hz). We then took a rare opportunity to record LFPs and spiking activity via microwires in V1 simultaneously. We found that RF sizes and temporal profiles measured from LGA and HGA closely matched those from spiking activity. In sum, this study reveals that spiking activity of neurons in human visual cortex could be well approximated by LGA and HGA in RF estimation and temporal profile measurement, implying the pivotal functions of LGA and HGA in early visual information processing.

## INTRODUCTION

Neurons in visual cortex initially encode the external world in a fragmental manner by restrictively responding to stimuli in a small portion of the visual field, namely the receptive field (RF) (Hubel and Wiesel, 1959). For decades, the RF concept, which interprets how external information is processed and represented internally, has served as a Rosetta stone of neuroscience. RF properties, such as size and location, are typically measured based on single-neuron activity. Since it is extremely rare to record single-neuron activity in humans due to technical difficulties and ethical considerations (Mamelak et al., 2014), our understanding of neuronal RF properties mainly derives from animal studies (Andoni et al., 2013). RFs in human visual cortex have been predominantly explored in functional magnetic resonance imaging (fMRI) studies using the population receptive field (pRF) method, which indirectly measures the collective RF of all active neurons in a voxel based on blood-oxygenation-level-dependent (BOLD) signals (Dumoulin and Wandell, 2008; Wandell et al., 2007; Wandell and Winawer, 2015), leaving the neuronal RF properties in humans nearly unknown. This long-overlooked barrier raises an intractable question: can neuronal RF properties in animals, even in non-human primates, be used to explain human visual perception (Spillmann and Werner, 1990)? Given accumulating evidence demonstrating anatomical and functional differences between human and non-human primate brains (Beaulieu-Laroche et al., 2018; Boldog et al., 2018; Eyal et al., 2016; Gabi et al., 2016; Kaas and Herculano-Houzel, 2017; Pryluk et al., 2019), it now becomes urgent to explore RF properties in human subjects through direct measurement of neuronal activities, such as local field potentials (LFPs) and spiking activity.

*In vivo* extracellular recording studies in human visual cortex are extremely rare. Using microwire recording in neurosurgery patients, two pioneering studies tried to map visual RFs based on spiking activity (Marg et al., 1968; Wilson et al., 1983). However, the precise locations of microwires were unclear due to technological limitations at that time. To the best of our knowledge, the only quantitative description of neuronal RFs estimated from multi-unit activities was obtained via two microwires implanted in V2/V3, which suggests that the RF sizes in humans were comparable to those in macaques (Self et al., 2016).

Most intracranial studies exploring RF properties in human visual cortex are based on LFPs recorded via macro-contacts (Bosking et al., 2017; Self et al., 2016; Winawer et al., 2013; Winawer & Parvizi, 2016; Yoshor, Bosking, Ghose, & Maunsell, 2006). LFP components are generated by neural circuits at various spatial scales and can be functionally heterogeneous in visual processing (Bartoli et al., 2019; Han et al., 2021; Hermes et al., 2014; van Kerkoerle et al., 2014). Therefore, RFs estimated from different LFP components could differ in size and location. For example, Winawer and colleagues (2013) found that RFs estimated from asynchronous and stimulus-locked LFP components slightly differed in size and location.

A fundamental question in neurophysiology is the relationship between LFPs and spiking activity. More specifically, can LFPs approximate spiking activity in RF estimation? This issue has been extensively investigated in animal studies (Belitski et al., 2008; Burns et al., 2010; Klink et al., 2021; Rasch et al., 2008; Ray et al., 2008a; Ray and Maunsell, 2011a; Ray and Maunsell, 2011b). In humans, the relationship between LFP components and spiking activity has been examined in auditory cortex (Mukamel et al., 2005; Mukamel et al., 2011; Nir et al., 2007; Zanos et al., 2012), motor cortex (Perge et al., 2014), and prefrontal cortex (Leszczyński et al., 2020). While some studies revealed a robust temporal correlation between spiking activity and high-frequency LFP components (Mukamel et al., 2005; Mukamel et al., 2011; Perge et al., 2014), others did not (Leszczyński et al., 2020; Nir et al., 2007; Zanos et al., 2012). Notably, this issue has never been examined in human visual cortex.

In the current study, we applied RF mapping procedures and intracranially recorded LFPs and spiking activity in human visual cortex (V1/V2/V3) to (1) quantify and compare neuronal RF properties estimated from different types of signals and (2) examine the temporal relationships between these signals. We first recorded LFPs via macro-contacts in 22 subjects undergoing invasive monitoring for the purpose of treating drug-resistant epilepsy (Experiments 1 and 2) (**Figure S1** and **Table S1**). Then, in 2 of the 22 subjects, we simultaneously recorded LFPs and spiking activity via microwires (Experiment 3) (**Table S2**). Based on a V1-V3 atlas (Benson et al., 2014) applied to the pre-implant T1-weighted MRI scan of individual subjects, we identified 244 macro-contacts localized in V1-V3 and 18 microwires in V1 (**Figure S1** and **Table S1**). LFPs were separated into three components: low-frequency activity (LFA, 0.5 to 30 Hz), low-gamma activity (LGA, 30 to 60 Hz), and high-gamma activity (HGA, 60 to 150 Hz) (Bartoli *et al.,* 2019; Parvizi and Kastner, 2018).

## RESULTS

### RFs estimated from LFA, LGA, and HGA

Experiment 1 aimed to compare locations and sizes of RFs estimated from LFA, LGA, and HGA. All subjects underwent a preliminary RF mapping procedure with 3° × 3° checkerboard stimuli (**Figure 1A**). To maintain fixation at the center of the screen, subjects were required to respond to color changes of the central fixation point throughout the experiment (response accuracy: 95.2 ± 0.9%, Mean ± SEM). Meanwhile, a 3° × 3° checkerboard was briefly presented at one of 99 (11 × 9 grid covering the full visual field) or 54 (6 × 9 grid covering the contralateral visual field) mapping positions (Yoshor *et al.,* 2007).

**Figure 1.**
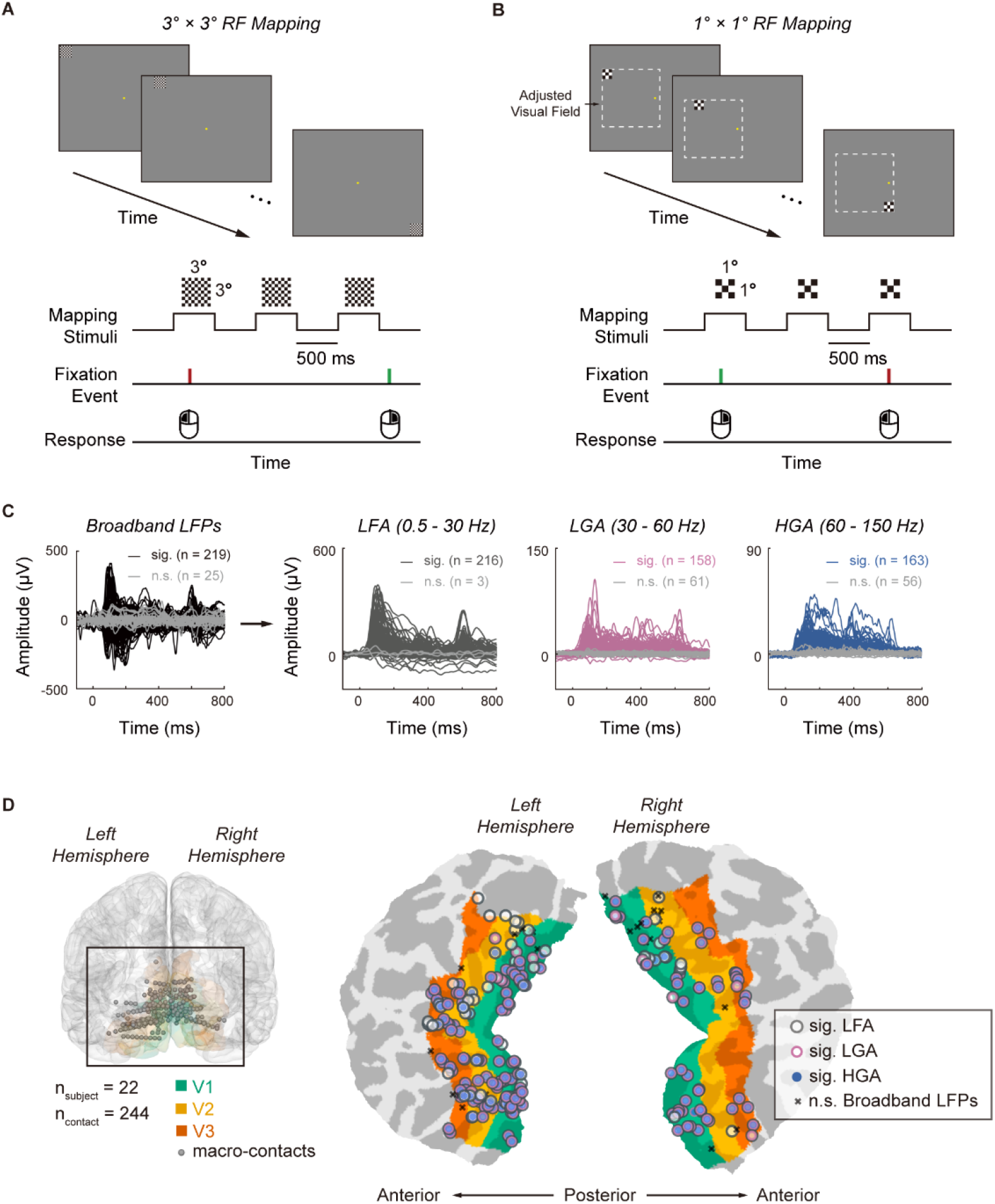
RF mapping procedures, visually evoked responses, and macro-contact locations. (A) Schematic description of the 3° × 3° RF mapping procedure used in Experiment 1. Subjects performed a fixation task while a 3° × 3° checkerboard stimulus was flashed at different positions on the monitor. (B) Schematic description of the 1° × 1° RF mapping procedure used in Experiments 2 and 3. Subjects performed a fixation task while a 1 ° × 1° checkerboard stimulus was flashed at different positions within an adjusted visual field *(dashed box),* which covered the responsive visual field in Experiment 1. (C) Visually evoked responses recorded from macro-contacts in Experiment 1. Waveforms of significant broadband LFPs *(black),* LFA *(dark grey),* LGA *(pink),* and HGA *(blue)* and non-significant waveforms (*light grey*) are shown. (D) Locations of macro-contacts visualized on a 3-dimensional template brain *(left)* and a flattened occipital patch *(right;* see **Figure S1** for locations of macro-contacts in individual brains). The colors on the brain indicate the retinotopic areas of interest *(green:* V1; *orange:* V2; *red*: V3). On the flattened occipital patch, the symbols indicate the significant LFP components identified for each macro-contact *(grey annulus:* LFA; *pink annulus:* LGA; *blue circle:* HGA). Macro-contacts without significant broadband LFPs were marked with *black* crosses. *Sig.:* significant; *n.s.*: non-significant.

We identified visually responsive macro-contacts by the presence of broadband visually evoked LFPs (0.5 – 200 Hz) to at least one mapping position (**Figure 1C**). Of all 244 macro-contacts in visual cortex, 219 (89.8 %) were visually responsive (V1: n_contact_ = 110; V2: n_contact_ = 66; V3: n_contact_ = 43). Next, we filtered the broadband LFPs into LFA, LGA, and HGA. 98.6% (216/219), 72.1% (158/219), and 74.4% (163/219) macro-contacts exhibited significant visually evoked LFA, LGA, and HGA, respectively (**Figures 1C, 1D**, and **Table S1**).

For each visually responsive macro-contact, we estimated RFs from LFA, LGA, and HGA (i.e., RF_LFA_, RF_LGA_, and RF_HGA_) separately, adopting a 2D-Gaussian fitting method described in previous ECoG studies (Nir et al., 2007; Yoshor et al., 2007). The size and location of an RF were defined as the averaged full width at half maximum (FWHM) of the two axes and the center of the fitted Gaussian function, respectively. RFs centered at or outside the border of the grids were excluded from further analyses. In total, we identified RF_LFA_, RF_LGA_, and RF_HGA_ for 81 macro-contacts (V1: n_contact_ = 41; V2: n_contact_ = 29; V3: n_contact_ = 11).

**Figure 2A** shows the estimations of RF_LFA_, RF_LGA_, and RF_HGA_ in an example V1 macro-contact (P440, macro-contact V01). We observed that the three RFs differed: the sizes of RF_LGA_ (size = 2.1°, location = [2.6°, −7.3°; azimuth, elevation] and RF_HGA_ (size = 2.2°, location = [2.6°, −6.3°]) were smaller than that of RF_LFA_ (size = 3.7°, location = [1.3°, −5.6°]). A one-way repeated-measures ANOVA found a significant main effect of LFP component (LFA/LGA/HGA; *F_2,160_* = 22.042, *p* = 3.506 × 10^-9^). Post hoc tests showed that the mean sizes of RF_LGA_ (2.2 ± 0.1°) and RF_HGA_ (2.4 ± 0.2°) were significantly smaller than that of RF_LFA_ (3.3 ± 0.2°; RF_LFA_ v.s. RF_LGA_: *p* = 1.289 × 10^-6^; RF_LFA_ v.s. RF_HGA_: *p* = 2.179 × 10^-6^, *Bonferroni* corrected; **Figures 2B** and **2C**). No significant size difference was found between RF_LGA_ and RF_HGA_ *(p* = 0.641). The RF sizes estimated from LFPs were within the range of previous estimations from LFPs in human subjects (Self et al., 2016; Yoshor et al., 2007). We then compared the locations of RF_LFA_, RF_LGA_, and RF_HGA_ in the polar coordinate system. One-way repeated-measures ANOVAs showed that there was no significant difference in either eccentricity (*F_2,160_* = 1.896, *p* = 0.153) or polar angle (*F_2,160_* = 0.954, *p* = 0.387).

**Figure 2.**
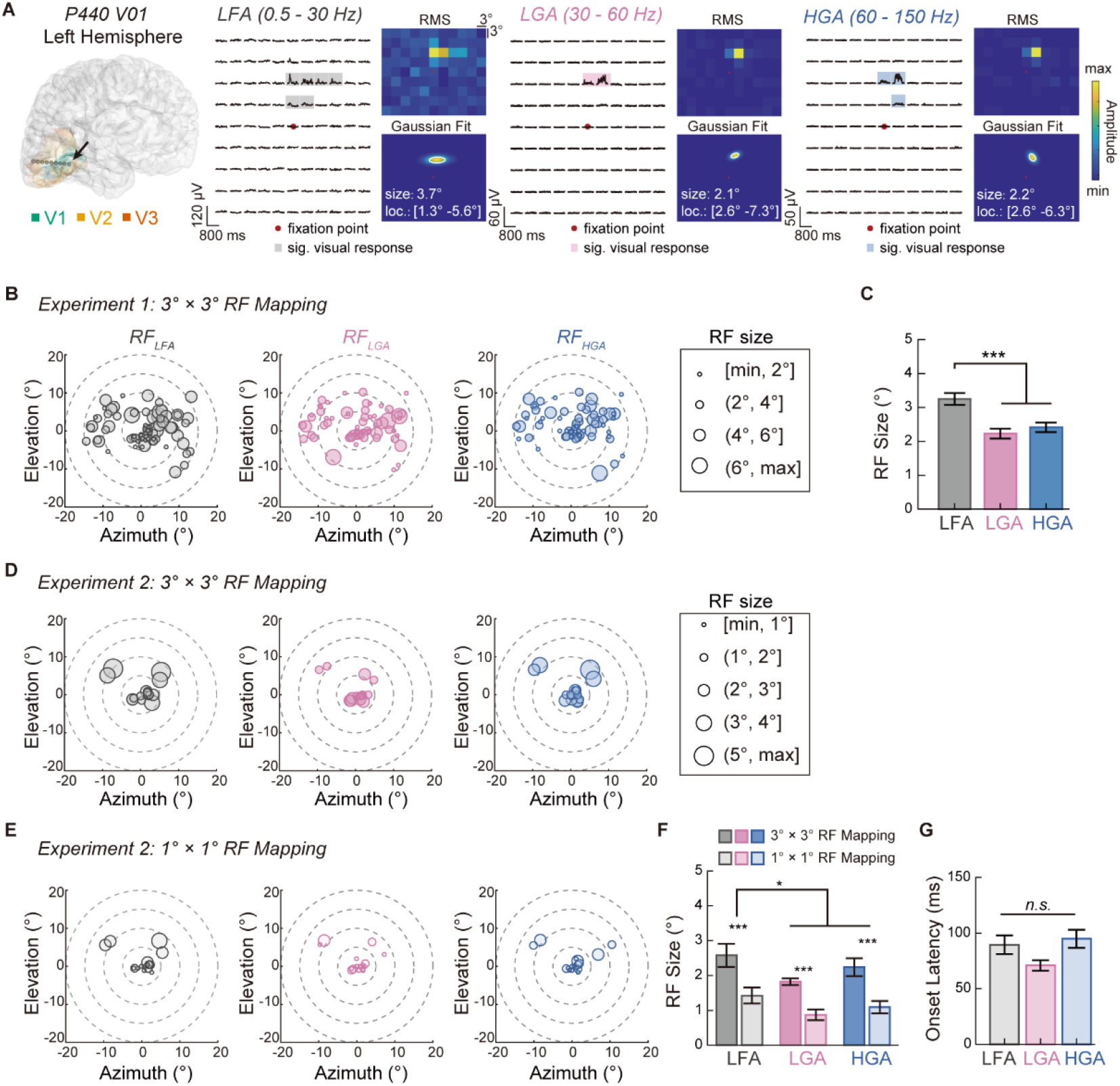
RF_LFA_, RF_LGA_, and RF_HGA_ mapping using macro-contacts. (A) Locations and sizes of the RF_LFA_, RF_LGA_, and RF_HGA_ estimated in an example macro-contact (subject P440, macro-contact V01). The stereo-electrode with macro-contacts is visualized in P440’s brain. For each LFP component, the averaged response waveforms to mapping positions *(red dot:* fixation point*; translucent areas:* significant responses), the RMS map, and the 2-D Gaussian fit of the RMS map are shown. The white ellipses show the RF contours. (B) Locations and sizes of RF_LFA_, RF_LGA_, and RF_HGA_ estimated with 3° × 3° checkerboard stimuli in Experiment 1. Each filled circle represents the RF of one macro-contact (n_contact_ = 81). The radius of the circles represents the size of the RFs. (C) Comparison of RF_LFA_, RF_LGA_, and RF_HGA_ sizes estimated in Experiment 1. (D) Locations and sizes of RF_LFA_, RF_LGA_, and RF_HGA_ estimated with 3° × 3° checkerboard stimuli in Experiment 2. (E) Locations and sizes of RF_LFA_, RF_LGA_, and RF_HGA_ estimated with 1° × 1° checkerboard stimuli in Experiment 2. (F) Comparison of RF_LFA_, RF_LGA_, and RF_HGA_ sizes estimated in Experiment 2. (G) Comparison of onset latencies among LFA, LGA, and HGA in Experiment 2. *RMS:* root mean square; *error bars:* standard error; * *p* < 0.05; ** *p* < 0.01; *** *p* < 0.001; *sig.:* significant; *n.s.:* none significant.

Although the 3° × 3° checkerboard stimuli used in Experiment 1 enabled us to identify RF locations quickly, their relatively large size inevitably led to an overestimation of RF sizes. Therefore, in Experiment 2, we performed a second mapping procedure with five subjects (n_contact_ = 17) using 1° × 1° checkerboard stimuli (i.e., 1° × 1° RF mapping; **Figure 1B**). We found that RF sizes estimated using 3° × 3° checkerboard stimuli (RF_LFA_: 2.6 ± 0.3°; RF_LGA_: 1.8 ± 0.1°; RF_HGA_: 2.2 ± 0.26°) were systematically larger than those estimated using 1° × 1° checkerboard stimuli (RF_LFA_: 1.4 ± 0.2°; RF_LGA_: 0.9 ± 0.2°; RF_HGA_: 1.1 ± 0.2°; **Figures 2D** and **2E**). A two-way repeated-measures ANOVA of Stimulus size (3° × 3°/1° × 1°) × LFP component (LFA/LGA/HGA) showed that both main effects were significant (Stimulus size: *F_1,16_* = 54.438, *p* = 1.558 × 10^-6^; LFP component: *F_2,32_* = 6.483, *p* = 0.004; **Figure 2F**). No significant interaction effect was found (*F_2,32_* = 0.415, *p* = 0.664). The sizes of RF_LGA_ and RF_HGA_ were still significantly smaller than those of RF_LFA_ (RF_LFA_ v.s. RF_LGA_: *p* = 0.046; RF_LFA_ v.s. RF_HGA_: *p* = 0.031; post hoc tests), while no significant difference was found between RF_LGA_ and RF_HGA_ *(p* = 0.223; **Figure 2F**), consistent with the findings in Experiment 1. We then compared RF locations using two-way repeated-measures ANOVAs (Stimulus size × LFP component). No significant main effect and interaction were found in either eccentricity (Stimulus size: *F_1,16_* = 0.097, *p* = 0.759; LFP component: *F_2,32_* = 0.330, *p* = 0.721; interaction: *F_2,32_* = 1.153, *p* = 0.328) or polar angle (Stimulus size: *F_1,16_* = 1.249, *p* = 0.280; LFP component: *F_2,32_* = 1.222, *p* = 0.307; interaction: *F_2,32_* = 0.021, *p* = 0.979). Together, these results demonstrate that the sizes of RF_LFA_ are remarkably larger than those of RF_LGA_ and RF_HGA_, while the locations of these three RFs are nearly identical.

We considered two possible explanations for the larger size of RF_LFA_. First, LFA may be modulated by feedback projections from neurons in higher visual cortex (Bastos et al., 2015; Jensen et al., 2015; Michalareas et al., 2016; van Kerkoerle et al., 2014), which have large RF sizes (Dumoulin and Wandell, 2008). If so, the latency of LFA should be longer than those of LGA and HGA. Second, LFA may reflect the lateral connectivity from neighboring neurons (Angelucci et al., 2017; Gilbert et al., 1990; Stettler et al., 2002), which would lead to similar latencies among LFA, LGA, and HGA. To test these two explanations, we compared the onset latencies of visually evoked LFA, LGA, and HGA at each RF center in Experiment 2. A repeated-measures ANOVA of the LFP component showed no significant main effect on latencies (LFA: 89.6 ± 8.3 ms; LGA: 71.0 ± 4.7 ms; HGA: 94.8 ± 8.1s ms; *F*_2,32_ = 2.662, *p* = 0.085), supporting the second explanation (**Figure 2G**).

### RF estimated from spiking activity

RF is typically defined by spiking activity. It remains elusive how RF defined by spiking activity relates to RF defined by LFP. Thus, in Experiment 3, we took a rare opportunity to simultaneously record LFPs and spiking activity using microwires implanted in V1 of two subjects (**Figures 3A** and **S2**).

**Figure 3.**
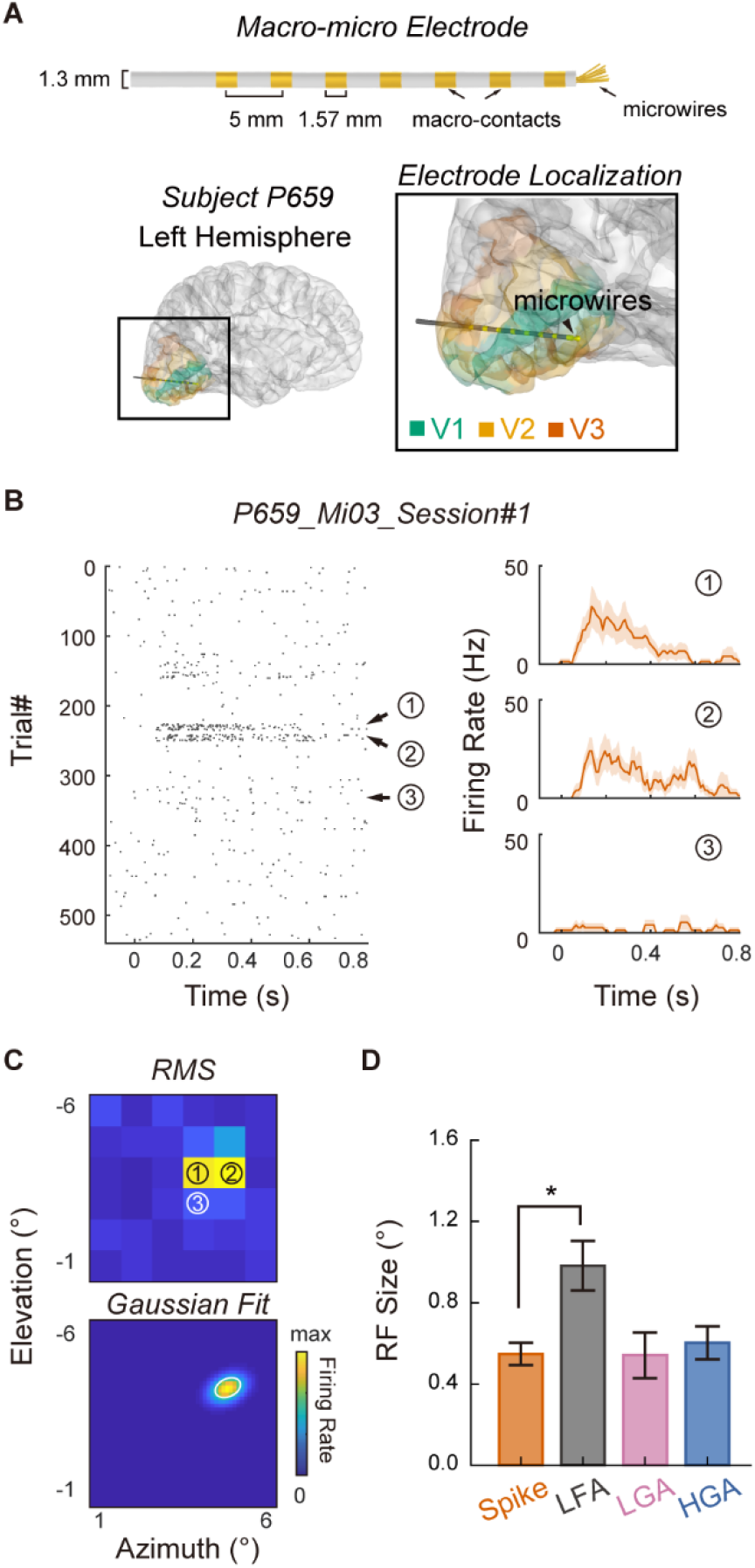
RF_spike_, RF_LFA_, RF_LGA_, and RF_HGA_ mapping using microwires. (A) Schematic description of the macro-micro electrode and the location of the microwires in P659. For P659, the 40 μm diameter microwires at the tip of the macro-micro electrode *(black* arrowhead) were localized in the ventral part of V1. (B) RF_spike_ mapping procedure of an example recording (P659_Mi03_Session#1). A raster plot of spikes around stimulus onset (sorted by mapping positions, *left*) and the averaged PSTH waveforms at three example positions (corresponding trials in the raster plot are marked by numbers, *right)* are shown. Shaded areas indicate the standard errors of the waveforms. (C) RMS map of spiking activity (*upper*) and two-dimensional Gaussian fit (*lower*) calculated for the example microwire shown in panel (B). The white ellipse indicates the RF_spike_ contour. *RMS:* root mean square. (D) Comparison of RF_spike_, RF_LFA_, RF_LGA_, and RF_HGA_ sizes (n_recording_ = 14). *Error bars:* standard error; * *p* < 0.05.

We isolated 55 units from two RF mapping sessions for P469 and four RF mapping sessions for P659 (**Table S2** and **Figure S3**). We identified 46 visually responsive units by the presence of significant visually evoked spiking activity to at least one mapping position. For microwires with visually responsive units, we further examined the significance of their visually evoked LFA, LGA, and HGA. For each mapping session, we defined a microwire recording with at least one visually responsive unit and visually responsive LFA, LGA, and HGA as a visually responsive recording (n_recording_ = 14, including six single-unit recordings and eight multi-unit recordings; **Table S2**).

For each visually responsive recording, we estimated RF sizes and locations from spiking activity (i.e., RF_spike_), LFA, LGA, and HGA (see **Figures 3B** and **3C** for an example recording). Note that RF_spike_ estimated for subject P469 had an average size of 0.5° at the eccentricity of about 4.3°, and RF_spike_ estimated for subject P659 had an average size of 0.6° at the eccentricity of about 6.2°. Previous studies have shown that the RF sizes of single-units in macaque V1 at similar eccentricities had a value of 0.1° – 1.5° (Blasdel and Fitzpatrick, 1984; Gattass et al., 1981; Levitt and Lund, 2002). Therefore, the RF_spike_ sizes in human V1 are generally in agreement with those in macaque V1.

We found a significant difference among the sizes of RF_spike_ (0.5 ± 0.1°), RF_LFA_ (1.0 ± 0.1°), RF_LGA_ (0.5 ± 0.1°), and RF_HGA_ (0.6 ± 0.1°) (*F_3,39_* = 6.078,*p* = 0.002; one-way repeated-measures ANOVA). Post hoc tests revealed that the sizes of RF_LFA_ were significantly larger than those of RF_spike_ *(p* = 0.036; **Figure 3D**). The sizes of RF_spike_ were not significantly different from those of RF_LGA_ and RF_HGA_ (both *p* > 0.05). To be noted, the sizes of RF_LFA_ were larger than those of RF_LGA_ and RF_HGA_ with marginal significance (RF_LFA_ vs. RF_LGA_: *p* = 0.060; RF_LFA_ vs. RF_HGA_: *p* = 0.085), consistent with the findings via macro-contacts in Experiments 1 and 2. For RF locations, no significant difference was found in either the eccentricity (*F_3,39_* = 0.347, *p* = 0.791) or the polar angle (*F_3,39_* = 2.619, *p* = 0.064; one-way repeated-measures ANOVAs).

We further quantified the spatial relationship between RF_spike_ and LFP RFs (RF_LFA_, RF_LGA_, and RF_HGA_) using an overlap coefficient (OC) index (Winawer and Parvizi, 2016). For two RFs, an OC index of 1 indicates perfect overlap, and 0 indicates no overlap (**Supplementary Methods** for details). Results showed that the three LFP RFs significantly differed in their OC indices with RF_spike_ (*F_2,26_* = 10.438, *p* = 4.701 × 10^-4^; repeated-measures ANOVA). Post hoc tests showed that, compared with RF_LFA_, RF_LGA_ and RF_HGA_ exhibited higher OC indices with RF_Spike_ (RF_LFA_ v.s. RF_LGA_: *p* = 0.033; RF_LFA_ v.s. RF_HGA_, *p* = 0.003; **Figure S4**). No significant difference was found between RF_LGA_ and RF_HGA_ *(p* > 0.1). These results suggest that RF_Spike_ can be better approximated by RF_LGA_ and RF_HGA_ than by RF_LFA_.

### Temporal relationships

The size similarity of RF_LGA_ and RF_HGA_ to RF_spike_ implies that they might have the same neuronal origin. One common evaluation of the neuronal origin of an LFP component is to calculate its temporal relationship with spiking activity (Mukamel et al., 2005; Ray et al., 2008a). In Experiment 3, we performed cross-correlation tests between the peri stimulus time histogram (PSTH) of spiking activity and the waveforms of LFA, LGA, and HGA evoked by the 1° × 1° checkerboard presented around each RF_spike_ center. We used the maximum cross-correlation coefficients to quantify the strength of LFP-spike correlations (i.e., LFA-spike, LGA-spike, and HGA-spike correlations).

As shown by the examples in **Figure 4A**, some recordings exhibited the strongest correlation between spiking activity and HGA, while others showed the strongest correlation between spiking activity and LGA. Low LFA-spike correlations were observed consistently across recordings. Overall, we found a significant main effect of the LFP-spike correlation type *(F_2,26_* = 12.329, *p* = 1.715 × 10^-4^; one-way repeated-measures ANOVA). Specifically, the LFA-spike correlation (0.380 ± 0.051) was significantly weaker than the LGA-spike (0.628 ± 0.034) and HGA-spike (0.686 ± 0.029) correlations (LFA vs. LGA: *p* = 0.023; LFA vs. HGA: *p* = 0.004; post hoc tests). No significant difference was found between the HGA-spike correlation and the LGA-spike correlation *(p* = 0.250; **Figure 4B**). These results revealed a close temporal relationship between LGA/HGA and spiking activity in human visual cortex.

**Figure 4.**
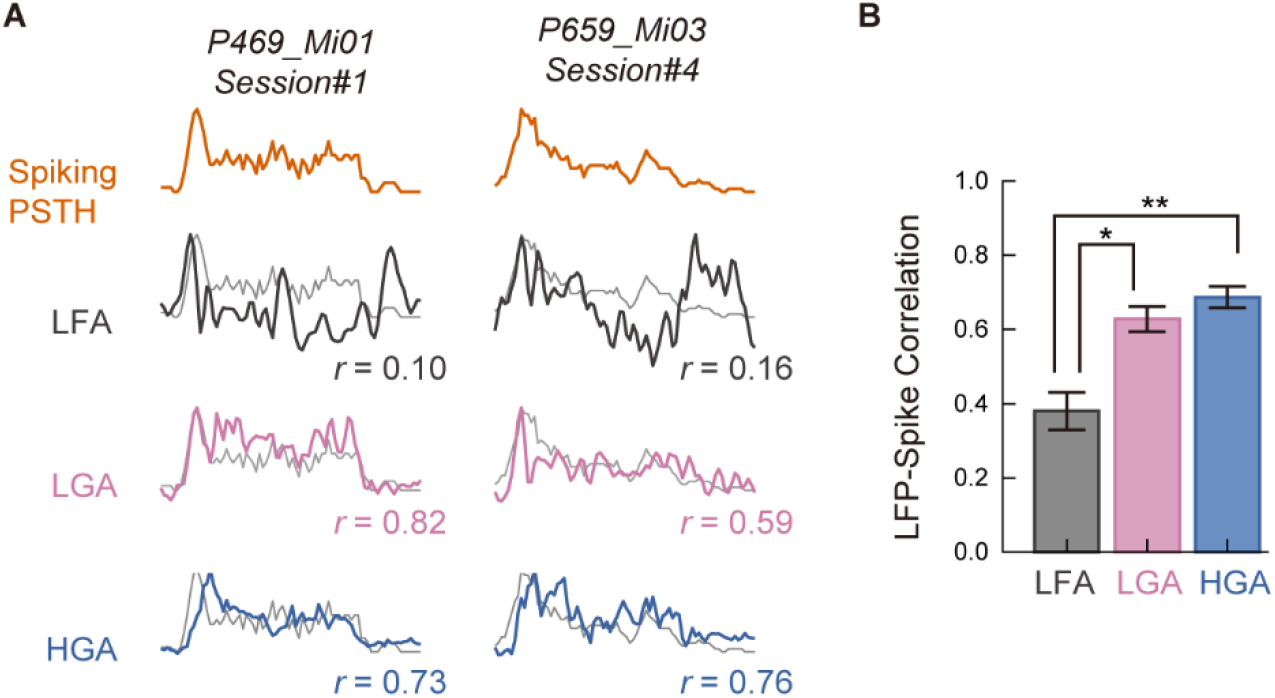
Temporal Correlations between spiking activity and LFP components. (A) Waveforms of spiking PSTH *(orange),* LFA *(dark grey),* LGA *(pink),* and HGA (*blue*) of two example microwires. The *grey* lines plotted with the LFP waveforms are the corresponding PSTH waveforms. The LFP-spike correlation coefficients (*r*) are also shown. (B) Comparison of the LFP-spike correlation coefficients among different LFP components (n_recording_ = 14). Compared with LFA, LGA and HGA showed stronger correlations with spiking activity in temporal profile. *Error bars:* standard error; * *p* < 0.05; ** *p* < 0.01.

## DISCUSSION

In this study, we estimated the sizes and locations of RF_LFA_, RF_LGA_, RF_HGA_, and RF_spike_, combining macro-contact and microwire recordings in human visual cortex. We found that although the four RFs were nearly identical in location, they differed in size. Specifically, the sizes of RF_LFA_ were remarkably larger than those of RF_LGA_, RF_HGA_, and RF_spike_, while no difference was found among the sizes of RF_LGA_, RF_HGA_, and RF_spike_. For the first time, we revealed that the RF_spike_ sizes in human V1 were generally consistent with those in macaque V1. Furthermore, LGA and HGA, but not LFA, showed high correlations with spiking activity in temporal profile. These findings suggest that, in terms of RF estimation and temporal profile measurement, RF_LGA_ and RF_HGA_ are potential surrogates of RF_spike_ if spiking activity cannot be obtained in human visual cortex.

We found that RF sizes estimated from LFA were significantly larger than those estimated from LGA, HGA, and spiking activity. Previous studies revealed that RFs in visual cortex have a center-surround organization (Bauer et al., 1995; Gieselmann and Thiele, 2008). That is, neurons respond to visual stimuli inside their RF and are also modulated by stimuli surrounding their RF (Allman et al., 1985; Angelucci et al., 2017). Thus, a possible explanation for larger RF_LFA_ sizes is that LFA is involved in surround modulation mechanisms, which integrate information inside and outside RF_spike_ and probably reflect dendritic computations of excitatory and inhibitory inputs (Adesnik et al., 2012; Buzsáki and Draguhn, 2004).

Gamma-range activities (> 30 Hz), which can be roughly divided into LGA and HGA, have been implicated in visual information processing (Brunet & Fries, 2019; Hermes et al., 2014; Hermes, Miller, Wandell, & Winawer, 2015; Lu et al., 2021; Peter et al., 2021; Zhang et al., 2020). There has been an ongoing debate on whether LGA and HGA in visual cortex are functionally distinct in encoding visual features. In monkey V1, LGA increased with stimulus size, but HGA decreased when the stimulus became larger (Dubey and Ray, 2020; Ray and Maunsell, 2011b). A recent study also revealed that LGA was prominently modulated by stimulus size and shape, while HGA was strongly modulated by location instead (Fischer and Wegener, 2021). In contrast, most intracranial recording studies in human visual cortex mainly investigated HGA (Davidesco et al., 2013; Golan et al., 2016; 2017; Martin et al., 2019; Zhou et al., 2019), leaving LGA largely unexplored. Until recently, Bartoli and colleagues (2019) found that LGA, but not HGA, was strongly tuned to contrast and hue, suggesting distinct functional roles of LGA and HGA in human visual cortex. In the current study, we found that the RFs estimated from LGA and HGA had similar sizes and locations in both macro-contact and microwire recordings. Given that LGA and HGA could be differentially modulated by stimulus size (Dubey and Ray, 2020; Ray and Maunsell, 2011b), one possible reason why we did not find the functional dissociation between LGA and HGA might be that we used relatively small and spatially discrete stimuli, whereas previous studies used larger and even full-screen visual stimuli (Bartoli et al., 2019; Hermes et al., 2015; Hermes et al., 2019; Winawer et al., 2013; Winawer and Parvizi, 2016).

Temporal correlation between gamma-range activities and spiking activity has been explored in animal models (Belitski et al., 2008; Burns et al., 2010; Rasch et al., 2008; Ray et al., 2008a; Ray et al., 2008b; Ray and Maunsell, 2011a; Ray and Maunsell, 2011b) and human subjects (Mukamel et al., 2005; Mukamel et al., 2011; Nir et al., 2007; Self et al., 2016). Using microwire recording in human auditory cortex, a series of studies revealed that the temporal profiles of gamma-range activities (40 to 130 Hz) were strongly correlated with spiking activity during both passive listening and resting state (Mukamel et al., 2005; Mukamel et al., 2011; Nir et al., 2007). To date, the only microwire study of human visual cortex also found that gamma-range activities (40-140 Hz) and neuronal spiking could be co-modulated by orientation and contrast (Self et al., 2016). In line with these findings, we revealed that the temporal profiles of both LGA and HGA were strongly correlated with neuronal spiking activities (mean correlation coefficients > 0.6) in human visual cortex. The strong HGA-spike correlation at the group level is also consistent with previous macaque studies (Kreiman et al., 2006; Liu and Newsome, 2006; Ray et al., 2008a; Ray and Maunsell, 2011b). Meanwhile, the strong spike-LGA correlation could be explained by a recently discovered class of neurons with strong LGA synchronization in macaque V1 (Onorato et al., 2020). In their study, Onorato and colleagues showed that these LGA-synchronized neurons were characterized by a narrow spike waveform. In the future, it would be of great interest to probe whether LGA and HGA in human visual cortex originate from different neuronal populations.

In sum, using direct electrophysiological recordings in human visual cortex, we explored the relationships between spiking activity and different LFP components (LFA, LGA, and HGA) in RF estimation and temporal profile measurement. We revealed that LGA and HGA, but not LFA, can closely approximate neuronal spiking activity in these two respects. Moreover, to the best of our knowledge, the current study is the first quantitative report of neuronal RF properties in human V1.

## EXPERIMENTAL MODELS AND SUBJECT DETAILS

### Human subjects

All human subjects were patients with drug-resistant epilepsy who underwent invasive stereo-electroencephalogram monitoring for potential surgical treatments at the Sanbo Brain Hospital of Capital Medical University (Beijing, China). LFPs were recorded from 22 subjects (17 males, mean age 27.3 years old) via macro-contacts (Experiments 1 and 2). Simultaneous recording of LFPs and spiking activity was performed with two subjects (P469: female, 32 years old; P659 male, 25 years old) via microwires (Experiment 3). Demographic information and implantation details are listed in **Table S1.** All subjects provided written, informed consent to participate in the experiments. All experimental procedures were approved by the Ethics Committee of the Sanbo Brain Hospital of Capital Medical University and the Human Subject Review Committee of Peking University.

## METHOD DETAILS

### Stereo-electrodes

Twenty patients were implanted with stereo-electrodes. Each stereo-electrode had 8-16 macro-contacts (0.8 mm in diameter, 2 mm in length, spacing 3.5 mm apart; Huake Hengsheng Medical Technology Co. Ltd., Beijing, China) (**Figure S1A**). All stereo-electrode implantations were determined based on clinical reasons.

### Macro-micro electrodes

Two patients (P469 and P659) were implanted with macro-micro electrodes in their visual cortex (BF08R-SP05X-000, WB09R-SP00X-014, AdTech Medical Instrument Corp., USA). Each macro-micro electrode had eight macro-contacts (1.3 mm in diameter, 1.57 mm in length, spacing 5 mm apart) and nine microwires at the tip (**Figure S1B**). All macro-micro electrode implantations were determined based on clinical reasons.

### Electrode localization and selection

For electrode localization, we first co-registered the post-implant CT images to the pre-implant T1-weighted MRI scans for each subject using the SPM12 toolbox (available at https://www.fil.ion.ucl.ac.uk/spm/software/spm12/; Penny et al., 2011). Then, we identified individual electrodes on the aligned CT images and calculated the coordinates of macro-contacts using the Brainstorm toolbox (available at http://neuroimage.usc.edu/brainstorm; Tadel et al., 2011). Since microwires were usually invisible on the post-implant CT images, their coordinates were estimated by combining the nearest macro-contact coordinates and the macro-micro electrode geometry (Bartoli et al., 2019; Self et al., 2016).

Only one subject (P469) participated in an fMRI retinotopic mapping experiment before electrode implantation. For P469, we defined her retinotopic visual areas (V1, V2, and V3) using a standard phase-encoded method (Engel et al., 1997; Sereno et al., 1995; for details, see **Supplementary Methods**, **Figures S2A** and **S2B**).

For the other 21 subjects, we selected macro-contacts localized in V1, V2, and V3 based on the anatomical identification in individual brains. We first performed cortical segmentation and reconstruction using individual pre-implant T1-weighted MRI scans in Freesurfer (version 6.0, Dale et al., 199). We then mapped V1, V2, and V3 onto individual cortical surfaces using a publicly available anatomical atlas (Benson et al., 2014) with codes from the Neuropythy toolbox (available at https://github.com/noahbenson/neuropythy/). We projected each macro-contact to the nearest vertex on the individual cortical surface using MATLAB (v2017b, MathWorks, MA, USA) function *dnsearch.* Macro-contacts were then assigned to V1, V2, and V3 based on the projected vertices. The *dnsearch* function also yielded the distance to the cortical surface for each macro-contact. Since the gray matter in human visual cortex is about 2-3 mm thick (Fischl and Dale, 2000) and macro-contacts had a maximum length of 2 mm, macro-contacts farther than 5 mm from the cortical surface were considered localized in white matter, and excluded from further analyses. Macro-contacts localized outside V1, V2, and V3 were also excluded from further analyses. For each macro-micro electrode, the anatomical identifications of microwires were referred to that of the nearest macro-contact.

To visualize all macro-contacts localized to visual cortex in a common space, we transformed the macro-contact coordinates into MNI coordinates and displayed them on a flattened cortical template *(cvs_avg35_inMNI152*) (**Figure 1C**). We also visualized the stereo-electrodes and macro-micro electrodes in individual brains (see **Figures S1** for locations of macro-contacts; see **Figures 3A** and **S2C** for locations of microwires).

### Experimental procedure

All experiments were conducted in quiet and dimly lighted patient rooms. Subjects were seated in bed during the experiments, with their head stabilized using a chin rest. Visual stimuli were generated and controlled using MATLAB (v2017b, MathWorks, MA, USA) and Psychtoolbox-3 extensions (Brainard & Vision, 1997; Kleiner, Brainard, & Pelli, 2007). They were presented on a laptop (14-inch, Thinkpad T590) or an LCD monitor (23.8-inch, Dell SE2416H) at a viewing distance of 40 to 60 cm.

#### Experiment 1: RF mapping using macro-contacts and 3° × 3° checkerboard stimuli

All 22 subjects participated in Experiment 1 (**Table S1**). We adopted an RF mapping procedure from a previous ECoG study (Yoshor et al., 2007). In each mapping run, a black-and-white checkerboard stimulus subtending 3° × 3° of visual angle was flashed at different mapping positions on the monitor to fill either an 11 × 9 (full visual field) or a 6 × 9 (half visual field contralateral to the implanted macro-contacts) grid (**Figure 1A**). Eight subjects received RF mapping of the full visual field, and 14 received RF mapping of the contralateral visual field. Each subject completed 15 to 20 mapping runs, resulting in 15 to 20 trials for each mapping position (one trial per mapping position per run). Checkerboard stimuli were presented for 500 ms at a temporal rate of 1 Hz. Subjects were asked to perform a fixation task by responding to color changes of the fixation point with mouse clicking (press the left button when changing to red and press the right button when changing to green). We tracked eye positions using an Eyelink Portable Duo tracker (SR Research, Canada) in 17 subjects (**Table S1**) and aborted trials in which the eye positions deviated away from the fixation point more than 1.0°. 13.9% of the total trials were excluded from further analyses.

#### Experiment 2: RF mapping using macro-contacts and 1° × 1° checkerboard stimuli

Five subjects participated in Experiment 2 (**Table S1**), which was similar to Experiment 1. For each run, a black-and-white checkerboard stimulus subtending 1° × 1° of visual angle was flashed at different positions on the monitor within an adjusted visual field optimized based on the results of Experiment 1. Each subject completed 15 to 20 mapping runs (one trial per mapping position per run). We tracked eye positions for all subjects. Trials in which the eye positions deviated away from the fixation point more than 1.0° were aborted (6.9 % of the total trials).

#### Experiment 3: RF mapping using microwires and 1° × 1° checkerboard stimuli

Two subjects (P469 and P659) participated in Experiment 3 (**Table S1**). The mapping procedure was identical to that in Experiment 2. To obtain sufficient single-unit and multi-unit signals, subjects completed multiple sessions (2 sessions for Subject P469, 4 sessions for Subject P659; **Table S2**). Each session consisted of 10 to 15 mapping runs (one trial per mapping position per run). Eye positions were monitored, and only 0.4% of the total trials were excluded using the same criteria in Experiments 1 and 2.

### Electrophysiological recording via macro-contacts

Macro-contact signals were recorded at a sampling rate of 512 Hz using a Nicolet video-EEG monitoring system (Thermo Nicolet Corp., USA) without any online filtering. Both the reference and ground electrodes were placed on the forehead of subjects. All further processing was performed offline.

### Electrophysiological recording via microwires

Microwire signals were recorded at a sampling rate of 32 kHz using an ATLAS neurophysiology system (Neuralynx Inc., USA). Signals were amplified using an HS-10-CHET pre-amplifier. For each macro-micro electrode, one microwire served as the local reference. We online monitored spiking activity using the Pegasus software (Neuralynx Inc., USA). Unfiltered raw signals were stored for offline extraction of both LFPs and spiking activity.

## QUANTIFICATION AND STATISTICAL ANALYSES

All signal processing and statistical tests were performed using publicly available toolboxes and custom scripts in Matlab (v2017b, MathWorks, MA, USA).

### Preprocessing of macro-contact signals

In Experiments 1 and 2, we imported raw macro-contact signals into the EEGLAB toolbox (version 14.1.1b; Delorme & Makeig, 2004) for visual inspection. Macro-contacts that contained epileptic activities or artifacts were excluded from further analyses. Signals were notch-filtered (50 Hz and harmonics) and then band-pass filtered (0.5 to 200 Hz) to generate broadband LFPs. Next, we filtered the broadband LFPs to obtain three LFP components: low-frequency activity (LFA, 0.5 to 30 Hz), low-gamma activity (LGA, 30 to 60 Hz), and high-gamma activity (HGA, 60 to 150 Hz). Two-way least-squares finite impulse response (FIR) filters were used *(eegfilt* function from the EEGLAB toolbox). We then extracted the amplitude envelope of each LFP component using the Hilbert transform. Finally, broadband LFPs and amplitude envelopes of the three LFP components were segmented around stimulus onset (−100 to 800 ms) and corrected against the baseline (−100 to 0 ms).

### Preprocessing of microwire signals

#### LFPs

Raw microwire signals in Experiment 3 were down-sampled to 2000 Hz for LFP extraction. We then obtained LFA, LGA, and HGA using the pipeline described above for processing macro-contact signals.

#### Spiking activity

In Experiment 3, we obtained spiking activity using the Osort toolbox (Rutishauser et al., 2006). We filtered the raw microwire signals with a zero-phase lag band-pass filter (300-3000 Hz; **Figure S3A**). Spikes were detected and sorted using an automatic algorithm (Rutishauser *et al.,* 2006; **Figures S3B** and **S3C**). We measured the quality of the isolated units using the following criteria: 1) the percentage of inter-spike intervals smaller than 3 ms (0.57 ± 0.09%); 2) the mean firing rate during each recording session (1.27 ± 0.18 Hz); 3) the signal-to-noise ratio (SNR) of the spike waveform (3.64 ± 0.15); 4) the modified coefficient of variation (CV2) (1.05 ± 0.01); 5) the pairwise distance between all pairs of isolated units on the same microwires (11.60 ± 0.61; **Figures S3D** to **S3H**) (Aquino et al., 2020; Fu et al., 2019; Kamiński et al., 2020; Minxha et al., 2020). Overall, we isolated 55 units from 6 recording sessions (7 units from subject P469 and 48 units from subject P659; **Table S2**).

For each unit, we segmented the spike trains into epochs from −100 to 800 ms around stimulus onset. Peri-stimulus time histograms (PSTHs) were constructed using non-overlapping 10-ms bins and then smoothed using a 50-ms square window with 10-ms steps.

### Identification of visually responsive macro-contacts and units

In Experiments 1 and 2, we defined visually responsive macro-contacts by the presence of a significant visually evoked broadband LFP to at least one mapping position. For a mapping position, the significance of a visually evoked broadband LFP was defined by two criteria: 1) the amplitude in the early response window (0 – 200 ms) exhibited at least one significant cluster longer than 10 ms (Wilcoxon rank-sum tests,*p* < 0.05; false discovery rate correction); 2) the standard deviation of the amplitude in the early response window was three times larger than the standard deviation of the baseline. For each visually responsive macro-contact, we tested the significance of LFA, LGA, and HGA using the same criteria described above.

In Experiment 3, we defined visually responsive units by the presence of significant visually driven spiking activity to at least one mapping position. For a mapping position, significant visually driven spiking activity was defined by two criteria: 1) the averaged firing rate at this mapping position was three standard deviations above the averaged firing rate at all mapping positions; 2) the standard deviation of firing rate in the early response window (0-200 ms) were three times larger than the standard deviation of the baseline. We also tested the significance of LFA, LGA, and HGA for microwires with at least one visually responsive unit using the same criteria described above for macro-contact signals. For each mapping session, we defined the recording from a microwire with both significant spiking activity and significant LFP components as a visually responsive recording.

### Estimation of the location and size of RFs

For each macro-contact or microwire recording, we estimated the size and location of the RF_LFA_, RF_LGA_, RF_HGA_, and RF_spike_ using the same pipeline. We calculated the root mean square (RMS) of the visually evoked response (LFA, LGA, HGA, or spiking activity) from 0 to 800 ms after stimulus onset at each mapping position. To reduce noise, we substituted the RMS value in mapping positions without significant neuronal responses with the averaged RMS value of all mapping positions. We fitted a two-dimensional (2D) Gaussian function to the RMS values for all mapping positions (Yoshor et al., 2007; Self et al., 2016). The RF location was defined as the coordinates of the fitted Gaussian function center. The RF size was determined by averaging the full width at half maximum (FWHM) of the two axes from the fitted Gaussian function.

### Evaluation of onset latency

In Experiment 2, we measured the onset latency of visually evoked LFA, LGA, and HGA using an established method (Bartoli et al., 2019; Foster et al., 2015). The response waveforms evoked by the checkerboard nearest the RF location were selected. We first smoothed single-trial responses with a 100-ms Gaussian sliding window. We then defined the response threshold as the 75^th^ percentile of the response amplitude across all trials for each macro-contact. For each trial, we marked the first time point at which the response amplitude exceeded the response threshold (lasted for at least 40 ms in the time window between −100 to 500 ms) and defined a 300-ms window of interest around the time point (100 ms before and 200 ms after). Next, we segmented the window into 50-ms bins with 90% overlap and fitted each bin using linear regression. The first time point of the bin with the largest slope and lowest residual error was then defined as the onset of the trial. The onset latency of a macro-contact was defined as the averaged latency across trials.

### Evaluation of temporal relationships

In Experiment 3, we used cross-correlation tests to examine the temporal correlations between spiking activity and LFP components (LFA, LGA, and HGA). For each visually responsive microwire, we selected the PSTH of spiking activity and waveforms of the three LFP components evoked by the 1° × 1° checkerboard presented around each RF_spike_ center. We then downsampled LFA, LGA, and HGA waveforms to match the 100 Hz sampling rate of the PSTH. Next, we calculated cross-correlation coefficients between the PSTH and the waveforms of LFA, LGA, and HGA with time lags between −100 to 100 ms. Finally, the maximal correlation coefficients were Fisher-*z* transformed for statistical analyses.

### Statistical analyses

We performed statistical analyses using custom scripts in Matlab (v2017b, MathWorks, MA, USA). To identify visually responsive LFPs, we used Wilcoxon rank-sum tests with false discovery rate (FDR) corrections for multiple comparisons. Moreover, we used repeated-measures ANOVAs to compare RF sizes, locations, latencies, and LFP-spike correlation coefficients. For post hoc tests, we reported Bonferroni corrected *p*-values. The alpha level was set at 0.05.

## Data and code availability

- The data that support the findings of this study are available from the Lead Contact upon request. The data are not publicly available because they could compromise research participant privacy and consent.
- Processed data and original codes used for analysis are available in a public repository as indicated in the key resources table (https://osf.io/evxq3/).
- Any additional information or code is available from the Lead Contact upon request.

## Acknowledgment

This work was supported by the National Science and Technology Innovation 2030 Major Program (2022ZD0204802, 2022ZD0204804) and the National Natural Science Foundation of China (31930053, 32171039) and Beijing Academy of Artificial Intelligence (BAAI).

## Author contributions

F.F., W.Q., and L.L. conceived the experiments. L.L., X.-F.W., J.-S.L., G.-P.C, and W.Q. collected the data. X.-F.W. and G.-M.L. performed the surgeries. L.L., G.-P.C, and W.Q. analyzed the data. W.L. provided technical assistance. F.F., W.Q., and L.L. wrote and revised the paper with input from all authors.

## Supplemental Information

### Supplementary Methods

#### fMRI Retinotopic Mapping

For subject P469 who was implanted with a macro-micro electrode, fMRI retinotopic mapping was conducted before implantation on a 3T Siemens Prisma MRI scanner at the Center for MRI Research at Peking University. We defined retinotopic visual areas of interest (V1, V2, and V3) using a standard phase-encoded method (Engel et al., 1997; He et al., 2019; Sereno et al., 1995) in which the subject viewed a rotating wedge and an expanding ring to create traveling waves of neural activity in visual cortex. MRI data were acquired using a 20-channel phase-array head coil. The visual stimuli were back-projected via a video projector (60 Hz refresh rate, 1024×768 spatial resolution) on a translucent screen inside the scanner bore. The subject viewed the visual stimuli through a mirror mounted on the head coil with a viewing distance of 73 cm. T1-weighted anatomical MRI scans were acquired before the retinotopic mapping session using a 3D-MPRAGE sequence (voxel size: 1 × 1 × 1 mm^3^), and the cortical surface was reconstructed using Freesurfer. Functional MRI scans were acquired using an echo-planner imaging (EPI) sequence (TE: 30 ms; TR: 2000 ms; flip angle: 90°; acquisition matrix size: 112 × 112; FOV: 224 × 224 mm^2^; slice thickness: 2.0 mm; gap: 0.3 mm; number of slices: 62; slice orientation: axial). Functional MRI scans were preprocessed, including slice timing correction, 3D motion correction, linear trend removal, coregistration with anatomical MRI scans, and spatial smoothing using the SPM12 toolbox in MATLAB. After preprocessing, the phase of BOLD signals was calculated using custom MATLAB codes and visualized using the TkSurfer tools in Freesurfer. Borders between visual areas were identified by the mirror reversals in the phase map. The retinotopic map indicated that the tip of the macro-micro electrode (co-located with the microwires) was localized in the ventral part of V1(**Figure S2**).

#### Quantification of RF Overlap

We quantified the spatial relationship between RF_spike_ and LFP RFs (RF_LFA_, RF_LGA_, and RF_HGA_) using a discrete version of the overlap coefficient (OC) index (Winawer and Parvizi, 2016). Specifically, for each LFP component, we calculated the number of mapping positions that showed either significant spiking activity (*N* [*spike*]) or significant LFPs (*N*[*LFP*]), as well as the number of mapping positions that showed both significant spiking activity and LFPs (*N* [*spike* ∩ *LFP*]). Then, we computed the OC as:

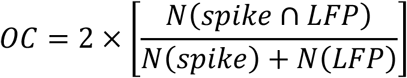

Therefore, an OC of 1 indicates complete spatial overlap between RF_spike_ and a certain type of LFP RF, and an OC of 0 indicates no overlap.

### Supplementary Tables

**Table S1.** Demographic and experimental information. Related to STAR Methods and Figure 1. Patient ID, age at the time of the experiment (years), gender (F = female, M = male), hemisphere of the electrodes (L = left, R = right), and experiment participation information (marked by *circles)* are listed. The numbers of macro-contacts with different selection criteria are listed (**Methods** for details).

| Patient ID | Age | Gender | Hemi-sphere | Participation in Experiments |  |  | #Contacts in Early Visual Cortex |  |  |  | #Visually Responsive Contacts |  |  |  | #Contacts with Significant LFA (0.5 - 30 Hz) |  |  |  | #Contacts with Significant LGA (30 - 60 Hz) |  |  |  | #Contacts with Significant LFA (60 - 150 Hz) |  |  |  |
| --- | --- | --- | --- | --- | --- | --- | --- | --- | --- | --- | --- | --- | --- | --- | --- | --- | --- | --- | --- | --- | --- | --- | --- | --- | --- | --- |
|  |  |  |  | Exp1 | Exp2 | Exp3 | Sum | V1 | V2 | V3 | Sum | V1 | V2 | V3 | Sum | V1 | V2 | V3 | Sum | V1 | V2 | V3 | Sum | V1 | V2 | V3 |
| 440 | 35 | M | L | ○ |  |  | 16 | 9 | 4 | 3 | 15 | 8 | 4 | 3 | 15 | 8 | 4 | 3 | 11 | 6 | 2 | 3 | 10 | 4 | 3 | 3 |
| 441 | 23 | F | L | ○ |  |  | 11 | 4 | 5 | 2 | 11 | 4 | 5 | 2 | 11 | 4 | 5 | 2 | 8 | 4 | 2 | 2 | 10 | 4 | 4 | 2 |
| 443 | 32 | M | L | ○ |  |  | 9 | 6 | 0 | 3 | 9 | 6 | 0 | 3 | 9 | 6 | 0 | 3 | 3 | 1 | 0 | 2 | 3 | 1 | 0 | 2 |
| 450 | 22 | M | L | ○ |  |  | 5 | 4 | 1 | 0 | 5 | 4 | 1 | 0 | 5 | 4 | 1 | 0 | 4 | 3 | 1 | 0 | 5 | 4 | 1 | 0 |
| 451 | 29 | M | L | ○ |  |  | 10 | 6 | 2 | 2 | 10 | 6 | 2 | 2 | 10 | 6 | 2 | 2 | 8 | 4 | 2 | 2 | 6 | 3 | 2 | 1 |
| 452 | 28 | M | L&R | ○ |  |  | 9 | 7 | 2 | 0 | 9 | 7 | 2 | 0 | 9 | 7 | 2 | 0 | 4 | 4 | 0 | 0 | 6 | 5 | 1 | 0 |
| 453 | 24 | M | L&R | ○ |  |  | 20 | 10 | 10 | 0 | 20 | 10 | 10 | 0 | 20 | 10 | 10 | 0 | 15 | 9 | 6 | 0 | 13 | 7 | 6 | 0 |
| 469 | 32 | F | L | ○ |  | ○ | 7 | 2 | 5 | 0 | 7 | 2 | 5 | 0 | 7 | 2 | 5 | 0 | 4 | 2 | 2 | 0 | 5 | 2 | 3 | 0 |
| 480 | 42 | M | L | ○ |  |  | 4 | 0 | 2 | 2 | 4 | 0 | 2 | 2 | 4 | 0 | 2 | 2 | 4 | 0 | 2 | 2 | 4 | 0 | 2 | 2 |
| 493 | 38 | F | L | ○ |  |  | 17 | 5 | 4 | 8 | 10 | 3 | 2 | 5 | 8 | 2 | 2 | 4 | 8 | 3 | 1 | 4 | 9 | 3 | 1 | 5 |
| 494 | 16 | F | L&R | ○ |  |  | 12 | 3 | 6 | 3 | 10 | 2 | 5 | 3 | 9 | 1 | 5 | 3 | 9 | 2 | 4 | 3 | 7 | 1 | 4 | 2 |
| 496 | 23 | M | L | ○ |  |  | 10 | 8 | 0 | 2 | 8 | 6 | 0 | 2 | 8 | 6 | 0 | 2 | 1 | 1 | 0 | 0 | 2 | 2 | 0 | 0 |
| 502 | 16 | M | L | ○ |  |  | 11 | 3 | 6 | 2 | 8 | 3 | 3 | 2 | 8 | 3 | 3 | 2 | 8 | 3 | 3 | 2 | 8 | 3 | 3 | 2 |
| 519 | 38 | M | R | ○ |  |  | 11 | 6 | 4 | 1 | 10 | 6 | 3 | 1 | 10 | 6 | 3 | 1 | 4 | 4 | 0 | 0 | 6 | 6 | 0 | 0 |
| 556 | 15 | M | L | ○ |  |  | 17 | 10 | 3 | 4 | 14 | 8 | 2 | 4 | 14 | 8 | 2 | 4 | 11 | 8 | 0 | 3 | 11 | 8 | 0 | 3 |
| 559 | 20 | M | L&R | ○ |  |  | 5 | 5 | 0 | 0 | 1 | 1 | 0 | 0 | 1 | 1 | 0 | 0 | 1 | 1 | 0 | 0 | 1 | 1 | 0 | 0 |
| 585 | 30 | M | R | ○ | ○ |  | 25 | 12 | 8 | 5 | 24 | 12 | 7 | 5 | 24 | 12 | 7 | 5 | 23 | 11 | 7 | 5 | 19 | 9 | 7 | 3 |
| 590 | 30 | F | L | ○ |  |  | 15 | 8 | 3 | 4 | 14 | 8 | 2 | 4 | 14 | 8 | 2 | 4 | 8 | 6 | 1 | 1 | 12 | 8 | 1 | 3 |
| 647 | 19 | M | R | ○ | ○ |  | 8 | 6 | 2 | 0 | 8 | 6 | 2 | 0 | 8 | 6 | 2 | 0 | 7 | 5 | 2 | 0 | 6 | 4 | 2 | 0 |
| 648 | 17 | M | L | ○ | ○ |  | 9 | 1 | 6 | 2 | 9 | 1 | 6 | 2 | 9 | 1 | 6 | 2 | 7 | 1 | 6 | 0 | 8 | 1 | 6 | 1 |
| 659 | 25 | M | L | ○ | ○ | ○ | 4 | 1 | 1 | 2 | 4 | 1 | 1 | 2 | 4 | 1 | 1 | 2 | 1 | 1 | 0 | 0 | 4 | 1 | 1 | 2 |
| 663 | 46 | M | L | ○ | ○ |  | 9 | 6 | 2 | 1 | 9 | 6 | 2 | 1 | 9 | 6 | 2 | 1 | 9 | 6 | 2 | 1 | 8 | 5 | 2 | 1 |
| #Sum |  |  |  | 22 | 5 | 2 | 244 | 122 | 76 | 46 | 219 | 110 | 66 | 43 | 216 | 108 | 66 | 42 | 158 | 85 | 43 | 30 | 163 | 82 | 49 | 32 |

**Table S2.** Mapping Sessions and Isolated Units in Experiments 3. Related to STAR Methods and Figure 3. P469 was tested with two sessions, and P659 was tested with four sessions. The numbers outside the parentheses are the number of isolated units for each microwire in each session. The numbers in the parentheses are the number of visually responsive units (see **STAR Methods**). The *asterisk* indicates that significant spiking activity, LFA, LGA, and HGA were identified (i.e., a visually responsive recording; n_recording_ = 14). In addition, we refer to recordings with one visually responsive unit as single-unit recordings (n_recording_ = 6) and those with more than one visually responsive unit as multi-unit recordings (n_recording_ = 8).

| Patient ID | Microwire Label | #Isolated Units (#Visually Responsive Units) |  |  |  |
| --- | --- | --- | --- | --- | --- |
|  |  | Session#1 | Session#2 | Session#3 | Session#4 |
| 469 | Mi01 | 1 (1)* | 1 (1)* | -- | -- |
|  | Mi02 | 1 (1)* | 2 (2)* | -- | -- |
|  | Mi07 | 0 (0) | 2 (2)* | -- | -- |
| 659 | Mi01 | 1 (1) | 2 (0) | 1 (1) | 1 (0) |
|  | Mi02 | 2 (2) | 1 (1)* | 2 (2) | 1 (1) |
|  | Mi03 | 2 (2)* | 2 (1)* | 2 (2)* | 2 (2)* |
|  | Mi04 | 2 (2)* | 2 (2) | 1 (1) | 2 (2) |
|  | Mi06 | 3 (3)* | 2 (2) | 2 (2)* | 2 (1)* |
|  | Mi07 | 2 (2) | 1 (1) | 1 (1) | 2 (0) |
|  | Mi08 | 2 (2) | 2 (1) | 1 (1) | 2 (0) |
| #Sum |  | 55 (46) |  |  |  |

### Supplementary Figures

**Supplementary Figure S1.**
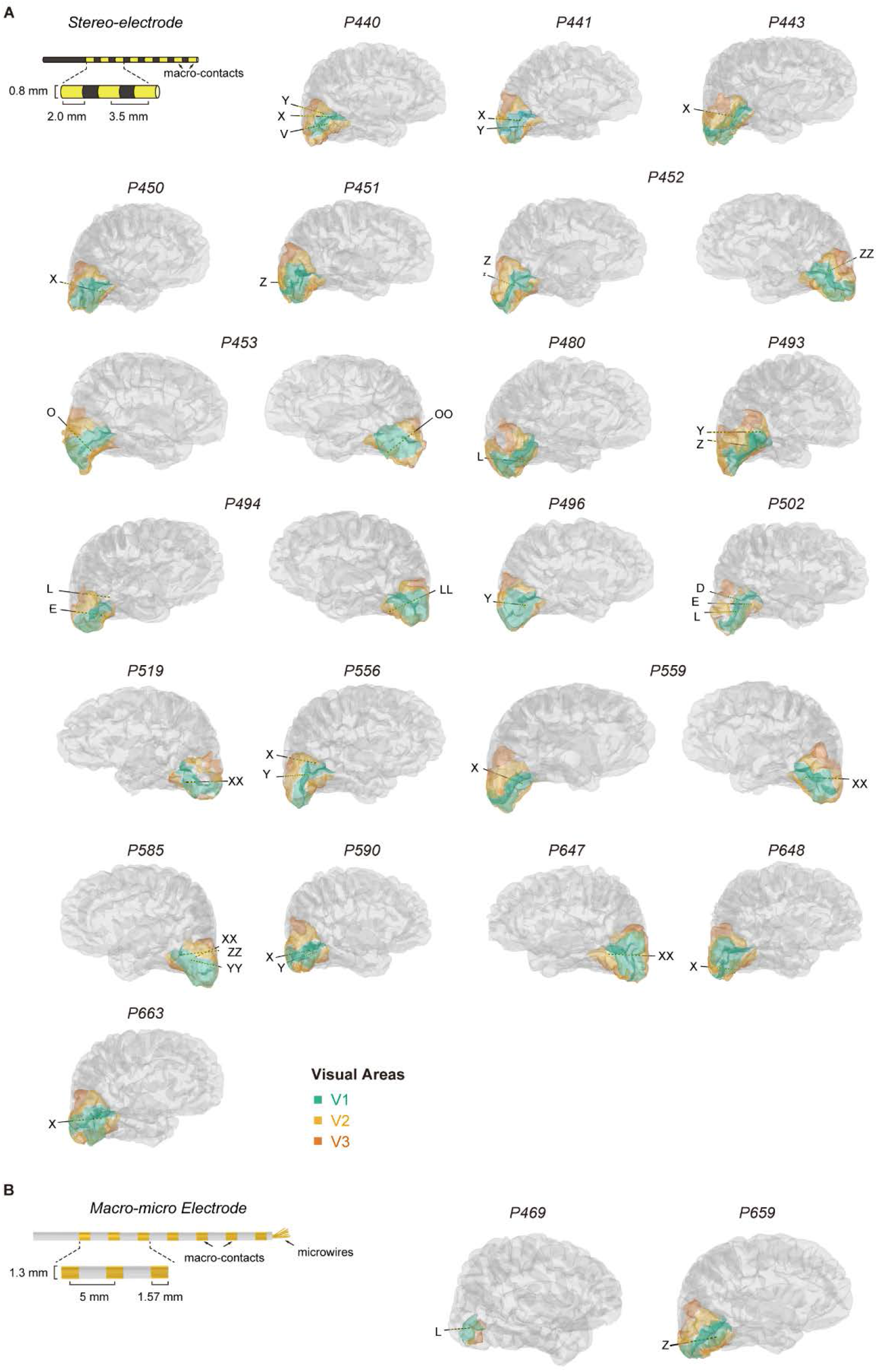
Individual cortical surfaces and locations of stereo-electrodes. STAR Methods and Figure 1. (A) Schematic description of an example stereo-electrode with 8 macro-contacts *(yellow)* and the locations of stereo-electrodes visualized in individual brains. The colors indicate retinotopic areas of interest (V1: *green;* V2: *orange;* V3: *red).* Only stereo-electrodes located in the occipital lobe are shown. (B) Schematic description of a macro-micro electrode and the locations of macro-micro electrodes visualized in individual brains. The colors indicate retinotopic areas of interest (V1: *green*; V2: *orange;* V3: *red).* In subject P469, the retinotopic areas of interest were defined by the fMRI retinotopic mapping procedure (see details in **Figure S2**).

**Supplementary Figure S2.**
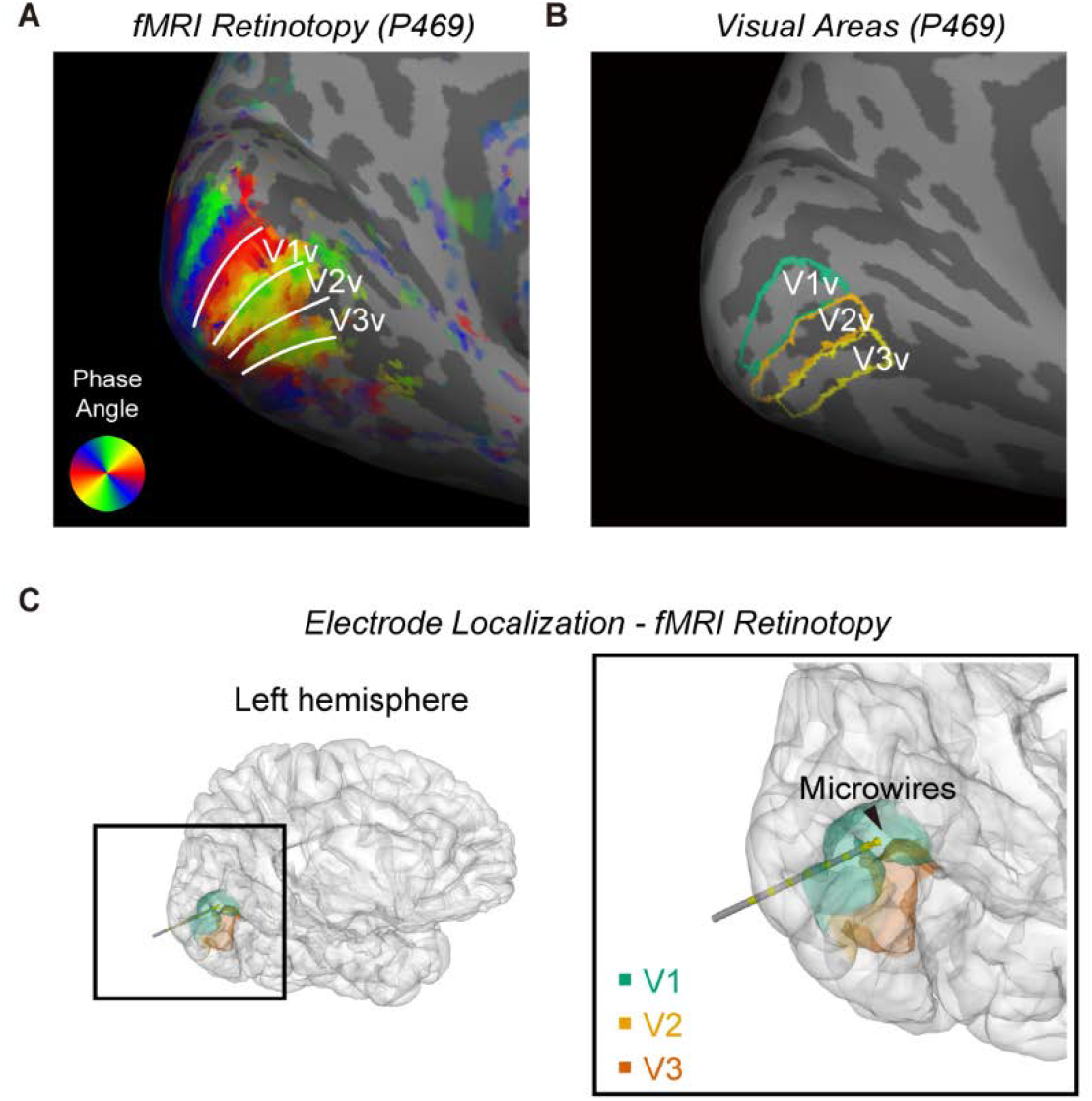
Retinotopic visual areas of subject P469 and location of microwires. Related to Figure 3. (A) Pre-implant fMRI measures of the retinotopic organization depicted on the inflated left hemisphere of subject P469. The white lines indicate the boundaries of ventral V1, V2, and V3 (V1v, V2v, and V3v, respectively). (B) Retinotopic areas of interest defined by the fMRI retinotopic mapping procedure. (C) Location of the macro-micro electrode visualized in the brain of subject P469. The tip of the macro-micro electrode (co-located with the microwires; *black* arrowhead) was located in ventral V1.

**Supplementary Figure S3.**
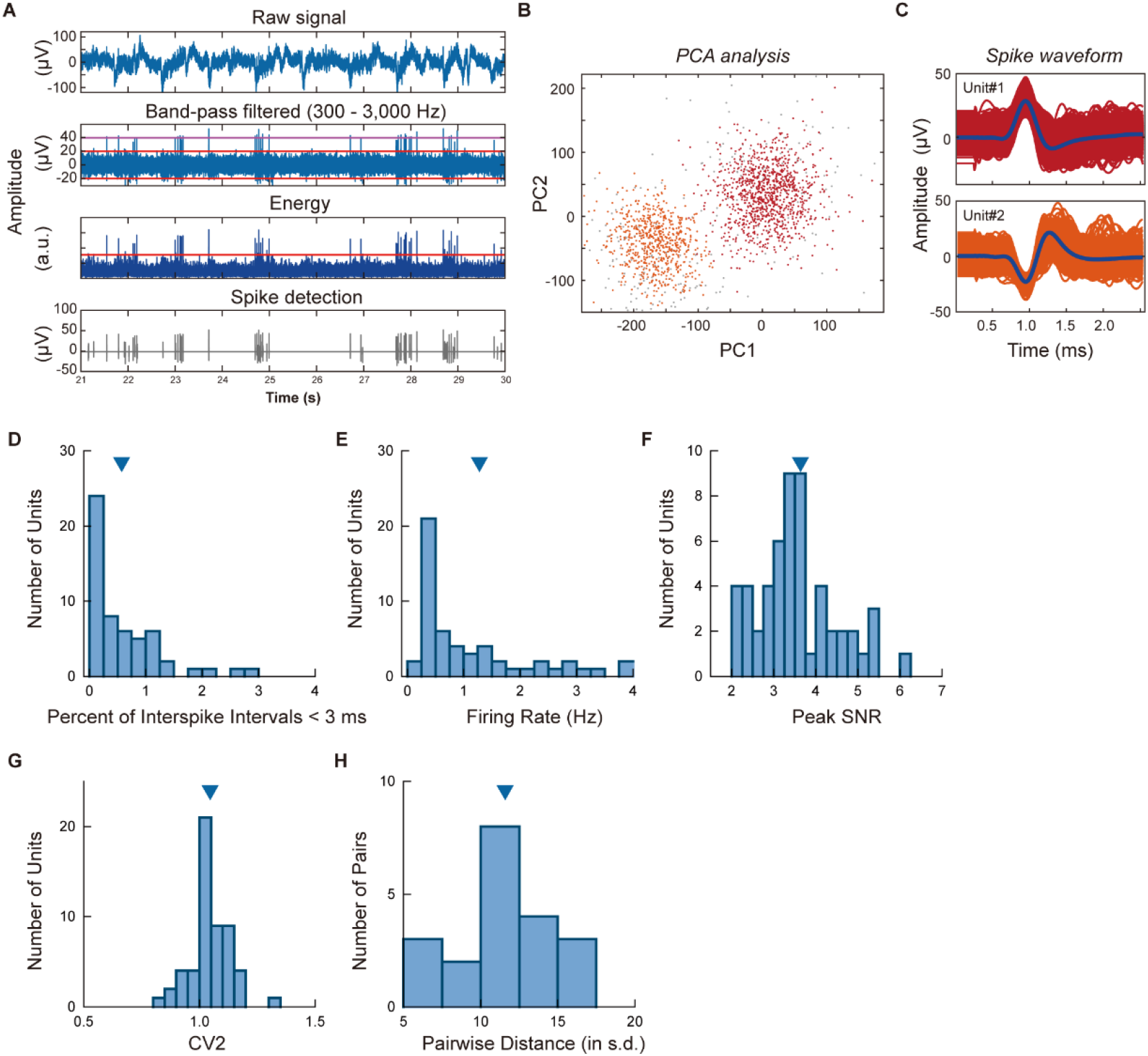
Spike detection and sorting. Related to STAR Methods. (A) Spike detection pipeline using the Osort toolbox. The raw signal *(top panel)* was band-pass filtered (300 to 3,000 Hz, *second panel; colored lines:* thresholds), and the local energy was calculated as the square root of the power using a running window of 1 ms *(third panel; colored line:* threshold). Spike detection was performed by thresholding the local energy *(bottom panel),* and spike waveforms were extracted from the band-pass filtered signal for sorting (*second panel; colored lines:* thresholds). (B) An example of the spike sorting results from microwire recording P659_Mi03_session#3. The post-hoc PCA plot of the first two principal components is shown, and two well-separated units could be observed (*red:* unit#1; *orange:* unit#2). (C) Waveforms of the two isolated units from P659_Mi03_session#3 *(blue:* mean spike waveform). (D) Histogram of the percentage of inter-spike intervals less than 3 ms for all isolated units (0.57 ± 0.09%; Mean ± SEM). (E) Histogram of the mean firing rate for all isolated units (1.27 ± 0.18 Hz). (F) Histogram of the peak SNR for all isolated units (3.64 ± 0.15). (G) Histogram of the CV2 value for all isolated units (1.05 ± 0.01). (H) Histogram of the pairwise distance for units isolated from the same microwire (11.60 ± 0.61). *Triangles*: mean value of the metric values.

**Supplementary Figure S4.**
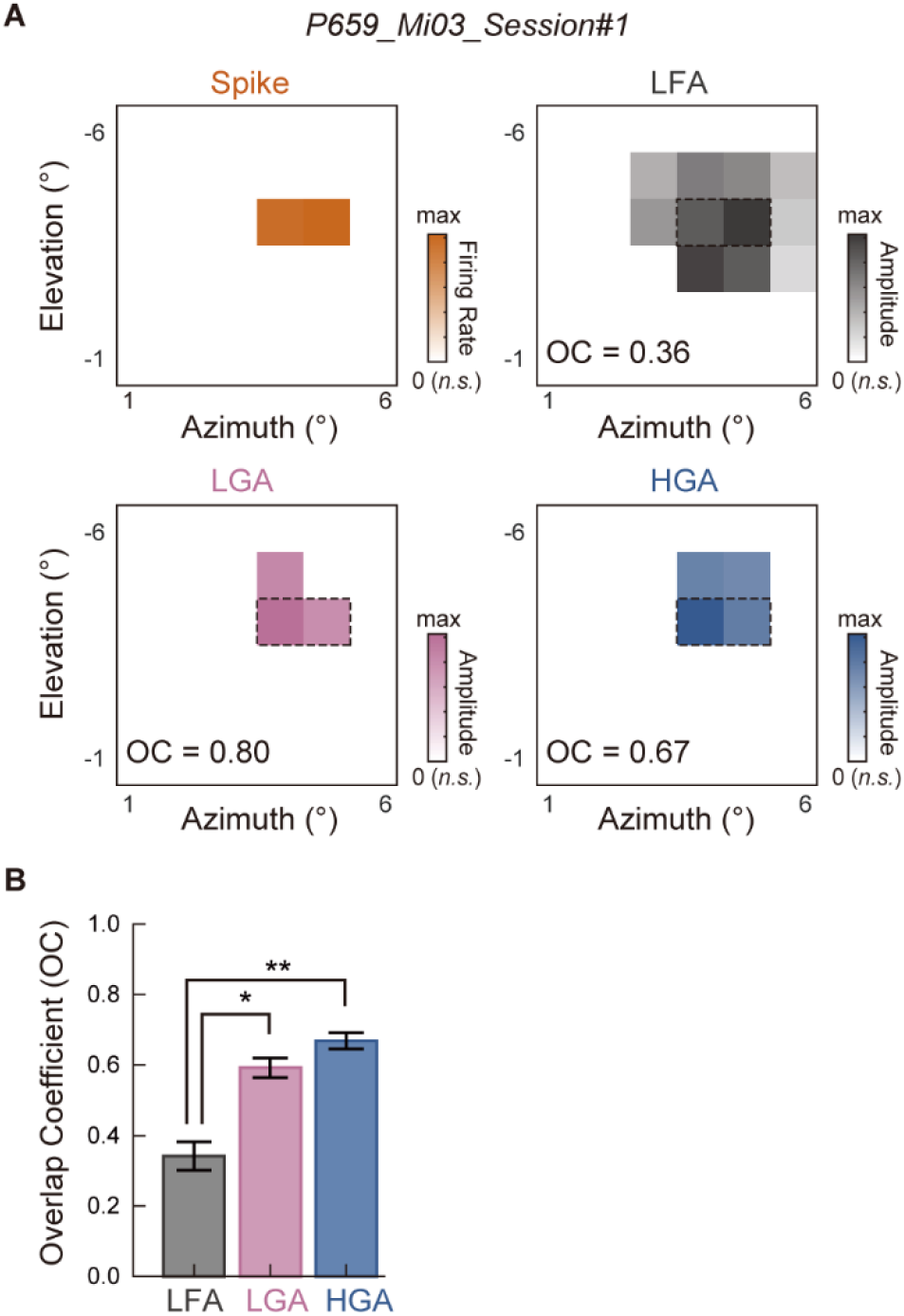
Overlap coefficients between LFP RFs and RF_spike_. Related to Figure 3. (A) Results of RF overlap coefficients (OCs) from an example recording (P659_Mi03_session#1). RMS maps are visualized for spiking activity, LFA, LGA, and HGA. RMS values were set to 0 for mapping positions without significant neural responses. In the RMS maps of LFPs, a *dashed* rectangle marks the mapping positions with significant visually evoked spiking activities. *RMS:* root mean square. (B) Comparison of overlap coefficients between RF_spike_ and each LFP RF. RF_LFA_, RF_LGA_, and RF_HGA_ significantly differed in their OC with RF_spike_ (RF_LFA_: 0.267 ± 0.062; RF_LGA_: 0.636 ± 0.090; RF_HGA_: 0.673 ± 0.060). Compared with RF_LFA_, RF_LGA_ and RF_HGA_ exhibited higher OCs with RF_spike_. *Error bars:* standard error; * *p* < 0.05; \*\**p* < 0.01.

